# DNA methylation database for gynecological cancer detection, classification and assay development

**DOI:** 10.1101/2024.07.01.601485

**Authors:** Joachim Boers, Ruben Boers, Jan Sakoltchik, Shatavisha Dasgupta, Lotte Martens, Kirke C.D. Tadema, Frederique Prevoo, Wilfred F.J. van IJcken, Henk van den Munckhof, Wim Quint, Heleen J. van Beekhuizen, Wim H. Quint, Folkert J. van Kemenade, Joost Gribnau

## Abstract

Changes in the genome wide DNA methylation landscape are hallmarks of cancer cells and precursor lesions of cancers. To capitalize on utilizing DNA methylation for detection and classification of cancer, we generated a DNA methylation database of gynecological cancers and associated healthy tissues using Methylated DNA sequencing (MeD-seq). We show that target cell enrichment to generate the database is crucial for marker discovery and report a wide range of novel biomarkers for classification and tissue of origin determination of gynecological cancers. We developed a subset of these novel biomarkers, both intragenic and intergenic, into a qMSP assays that detect all gynecological cancers at once or specific gynecological cancer subtypes, as well as cancers that are not part of our database. The database generated in this study not only provides the foundation for cancer detection, classification and biomarker discovery, but also for treatment monitoring of cancers using MeD-seq on liquid biopsies.

## Introduction

Cancer is among the leading causes of mortality worldwide and is generally associated with a poor prognosis due to advanced disease progression prior to presentation. It is challenging to improve prognosis as early stages of the disease tend to be asymptomatic and therapeutic windows for most anticancer treatments are generally narrow. Gynecological cancers are among the group of malignancies with a growing annual worldwide incidence, and are expected to account for 2.2 million yearly cases worldwide in the year 2050 (https://gco.iarc.who.int/). Many gynecological cancers are detected in an advanced disease state and as a result are associated with increased morbidity, decreased treatment options and a high mortality. Currently, early detection of gynecological cancers is limited to HPV-based cervical cancer screening programs. However, HPV screening is a risk-based test which requires further triage and its applicability is limited to HPV induced cancers and is not a reliable method for the detection of endometrial and ovarian cancers. Therefore, a more comprehensive screening test is needed that can be applied to detect the majority of gynecological cancers, including HPV-independent cancers.

Cancer development and progression is associated with, and/or directed by extensive epigenomic changes including changes in DNA methylation. DNA methylation plays a crucial role in gene regulation and occurs mainly at CG dinucleotide sequences in mammalian genomes [1]. In HPV-induced cancers of the cervix and vulva HPV infection is the key driver of DNA methylation changes as the HPV genes E6 and E7 affect the activity of DNA methyltransferases [2]. For HPV-independent cancers DNA methylation changes are often induced by mutations in genes involved in epigenetic processes such as DNA methyltransferases [3]. Consequently, single events such as infection or a mutation can result in massive genome-wide changes in DNA methylation providing a wide range of targets to detect and classify cancer, to study cancer progression or for applications in personalized medicine.

Previous studies have shown that malignant cells are present in cervical swabs and prove that it is possible to define cancer-specific DNA methylation markers in these samples using targeted approaches such as methylation sensitive and/or dependent PCRs [4–7]. Similarly, targeted assays have been applied to detect cancer-associated DNA methylation changes in cell-free DNA (cfDNA) obtained from blood [8]. The vast majority of DNA methylation marker regions used in currently available assays were found by limited genome wide DNA methylation profiling using methylation arrays [9, 10] in studies focusing on human cell lines [11, 12] or other types of cancer, and therefore potentially fail to detect more sensitive regions that could be used to detect the cancer type of interest [13]. Methylation arrays use pre-designed probes that only interrogate a small subset (2-4%) of all CG dinucleotides in the human genome. This limits the search for DNA methylation biomarkers, potentially missing powerful biomarkers. In addition, because current targeted DNA methylation assays are validated on cervical cancer and its precancerous lesions, their ability to detect other gynecological cancers remains unclear. Furthermore, current assays rely on a single or a few marker regions, whereas genome wide approaches are preferred for cancers with a more heterogeneous DNA methylation spectrum and highly sensitive applications such as disease monitoring or determination of the minimal residual disease (MRD).

In recent years we have developed and optimized Methylated DNA-sequencing (MeD-seq) to generate genome-wide DNA methylation profiles of healthy and diseased tissue [14]. MeD-seq utilizes a DNA methylation dependent restriction enzyme to generate 32 base-pair methylated fragments, that are isolated and sequenced. MeD-seq detects 50% of all potentially methylated CpG dinucleotides genome-wide, and can be applied on very small quantities of DNA isolated from Formalin-Fixed Paraffin-Embedded (FFPE) material. Here, we applied MeD-seq to generate genome-wide DNA methylation profiles of the majority of gynecological cancers and associated healthy tissues using Laser Capture Microdissection (LCM) for tumor enrichment. This DNA methylation atlas was used to define marker regions able to detect DNA methylation changes present in all gynecological cancers, subgroups of gynecological cancers or marker regions specific for one gynecological cancer type. We developed quantitative Methylation-Specific PCR (qMSP) assays for the detection of all gynecological cancers at once in biopsies using four different marker regions. We validated our assay to detect (pre-)stages of cervical cancer on cervical swab material from a multicenter study (EVAH) which consist of women with an abnormal Pap smear result who were referred for colposcopic evaluation.

Our studies indicate that LCM-mediated tumor enrichment is crucial for selection of the most predictive marker regions using MeD-seq for DNA methylation profiling. We also show that our DNA methylation database of gynecological cancers and controls provides numerous differentially methylated regions (DMR’s) that can be applied as cancer-specific or general multi-cancer markers, and that identified marker regions can be easily translated into a qMSP assay. In addition, our atlas shows potential for molecular classification of precursor lesions in cervical cancer. We believe that both genome wide DNA methylation profiling by MeD-seq and our localized qMSP assay can be used for cancer detection, classification and monitoring treatment outcome. In addition, the qMSP assay shows promising potential for the use in population-based screening for gynecological cancers, including cervical cancer.

## Results

### Tumor enrichment is crucial for genome wide DNA methylation profiling

MeD-seq is a sensitive technology that can be applied on very small quantities of DNA (5 ng) facilitating the analysis of FFPE tumor material through LCM (Fig. 1a). First, we tested whether enrichment of tumor material through LCM generates more informative DNA methylation profiles for discovery studies. We compared DNA methylation profiles of cervical carcinomas obtained through LCM of pure tumor regions with DNA methylation profiles obtained from sections of the same cervical carcinomas with varying loads of tumor cells and surrounding normal cells. We selected 10 whole tumor sections (WTS), two of which had 100% tumor contribution and eight with tumor contribution ranging from 70% to as little as 25%, and from these eight samples cells were collected both with and without LCM tumor enrichment (Fig. 1b). We established genome wide DNA methylation profiles using MeD-seq and compared the number of DMRs in cancer cells compared to cervical controls for both WTS and LCM-isolated samples. This analysis revealed a significantly higher number of DMRs in the LCM-enriched samples (Fig. 1c). In addition, DMRs observed with LCM enriched samples displayed an increased fold change per DMR compared to those obtained with WTS (Fig 1d). Hierarchical clustering based on WTS or LCM DMRs showing a >2-fold change separated all cancers from controls and showed that nearly all identified DMR’s are hypermethylated in cervical carcinomas (Fig. 1e,f). Interestingly, hierarchical clustering of WTS, LCM and control samples, reveals that WTS samples with low tumor contribution can still be distinguished from healthy tissue samples but do cluster separate from samples with a 100% tumor contribution (Fig. 1f). Therefore, once DMRs between cancer and healthy tissues are detected using highly tumor enriched reference sample sets, additional samples do not require a high tumor contribution in order to detect cancer. We conclude that tumor enrichment is crucial to establish genome wide DNA methylation profiles to detect novel DMRs.

**Figure 1.**
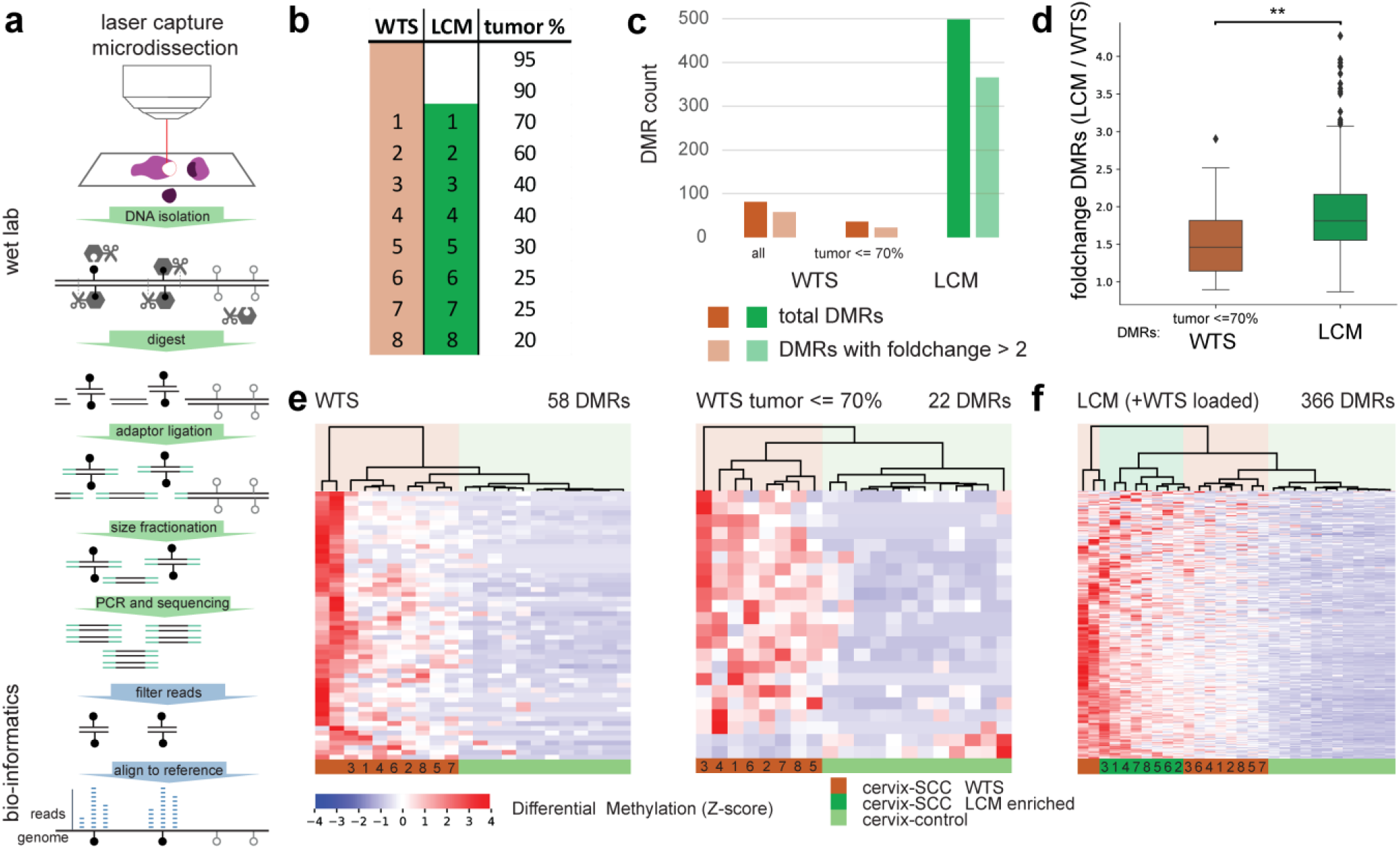
MeD-seq workflow and tumor enrichment. (a) MeD-seq workflow, tumor cells were enriched using LCM, and DNA was subsequently isolated from enriched tumor material and healthy control material. DNA is digested by a methylation dependent restriction enzyme and sequencing adaptors are ligated to 32 bp restriction fragments followed by size selection and sequencing. Sequenced reads are filtered based on restriction site position and selected reads are mapped to the human genome. (b) Overview of WTS cancer samples with associated tumor percentages. Eight WTS samples with tumor percentage <=70% were LCM enriched. (c) DMR detection comparing cancer to controls, all WTS, only WTS tumor <=70% and only LCM samples. (d) Foldchange of DMRs observed using WTS<=70% and LCM enriched samples, y-axis shows DMR-foldchange between average LCM sample read count divided by average WTS (<=70%) read count. Outliers are displayed as black diamonds. ** p < 0.01 (e) Heatmap based on WTS DMRs, hierarchical-clustered samples; WTS cervical carcinomas (brown) and healthy controls (light green) (z-score per DMR is shown). All WTS samples (left), WTS <=70% (right). WTS samples which were also analyzed by LCM are numbered according to figure 1b. (f) Heatmap based on LCM DMRs, hierarchical-clustered samples; WTS cervical carcinomas (brown), healthy controls (light green) and LCM (dark green) samples are indicated by corresponding numbers from figure 1b.

### A DNA methylation atlas of gynecological cancers and its origin

To identify DNA methylation changes associated with cancer development and generate a comprehensive DNA methylation atlas of the female genital tract, we combined MeD-seq and LCM to acquire DNA methylation profiles from various gynecological cancers and their associated healthy tissues (vulva, cervix, endometrium, fallopian tube, fimbriae and ovary). Our previous studies indicated an inter-patient variability associated with SNPs and copy number variations. To establish the minimum number of required samples, we performed bootstrapping analysis based on detection of DMRs between an increasing number of individual cervix squamous cell carcinoma (SCC) cancers indicating that the number of DMRs stabilized with a sample size above 7 (Supp. Fig. 1a). We therefore aimed for 10 samples per reference sample group, and, despite some rare cancers not being available in sufficient quantities, we succeeded for the majority of gynecological cancers (Fig 2a).

**Figure 2.**
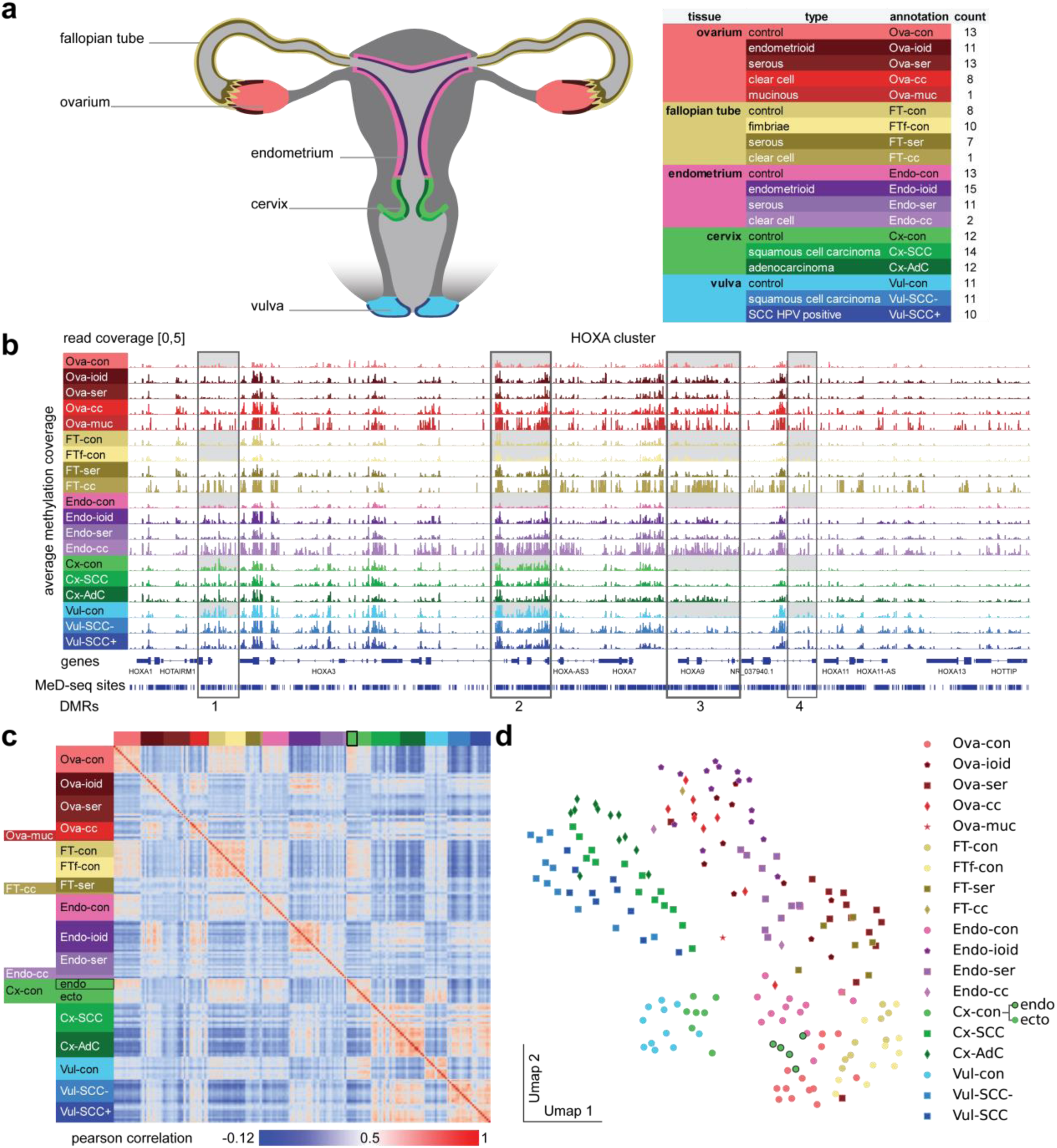
DNA methylation atlas of gynecological cancers and associated tissue origin. (a) Overview of the anatomical location where samples were collected and their abbreviations. Number of included samples per tissue type is shown right. (b) MeD-seq data at the HOXA gene cluster visualized in a genome browser (IGV). Shown are average methylation levels per sample group. Tissue-specific DNA methylation differences present in healthy tissues are highlighted in box 1 to 4 with a gray background. (c) Pearson correlation between samples using promoter, gene body and CpG island regions displaying variability in DNA methylation. Cervix controls are separated into endocervix and ectocervix groups. (d) UMAP based on DNA methylation of promoter, gene body and CpG island regions (same as in 2c). Endocervix (black outline) and ectocervix (no outline) separation of cervical control samples.

Examination of DNA methylation in the HOXA cluster, known to be differentially methylated during development in different embryonic lineages, confirms differences between healthy control tissues reminiscent of their distinct embryonic origin (Fig. 2b). To compare the DNA methylation landscape between the different cancers and controls we quantified the MeD-seq read counts for each CpG-island (genomic regions enriched for CG dinucleotides), gene promoter and gene body per individual sample and established the Pearson correlation coefficient in comparisons between all samples and between cancer and control types using the average DNA methylation profile per sample group. This analysis revealed a clear separation of all cancer types and controls, highlighting the unique changes associated with each individual cancer (Supp. Fig. 1b). In addition, comparison of DNA methylation profiles in between individuals with the same cancer types indicated that most cancer associated changes are relatively homogeneous within cancer types (Fig. 2c). The exception were the serous cancers (from ovary, fallopian tube and endometrium) which showed much less correlation in relation to each other and within their own group. Visualization of cancer and control samples based on uniform manifold approximation and projection of dimension reduction (UMAP) based on all observed DNA methylation changes confirms the clear distinction between cancers and controls and between several cancer types that cluster as separate groups (Fig. 2d). Interestingly, this UMAP revealed two distinct groups of cervix controls, and indeed, subsequent pathological examination of these samples indicated that one group, which clustered close to the vulva controls, consists of stratified epithelium (ecto cervix), whereas the other group, that clustered with endometrium, consisted of columnar epithelium (endo cervix).

### HPV detection and genotype classification using MeD-seq

Development of several gynecological cancers is driven by HPV infections. Persistent expression of high-risk HPV sub-types, particular HPV16 and HPV18, is responsible for most cervix SCCs, the majority (85%) of cervix adenocarcinomas (ADCs), and a substantial portion of vulva SCCs (30%). Continued expression of viral proteins leads to transformation of epithelial cells, via intermediate steps to cancer. Persistent HPV infection eventually leads to integration of HPV into the genome and this integrated viral DNA can become methylated [15]. To test whether MeD-seq is able to detect methylated viral sequences, we added the different HPV viral genomes as reference for quantification of MeD-seq reads (Fig 3a,b). SPF10 PCR-mediated detection of HPV subtypes confirmed the presence of high-risk viral genomes (HPV16 and HPV18) in nearly all HPV-induced cancer samples (Fig. 3c). Analysis of our MeD-seq reads revealed a similar picture and correctly detected DNA methylation of HPV virus associated sequences in 92% of the cervix AdCs, and 93% and 90% of the cervix and vulva SCCs respectively.

**Figure 3.**
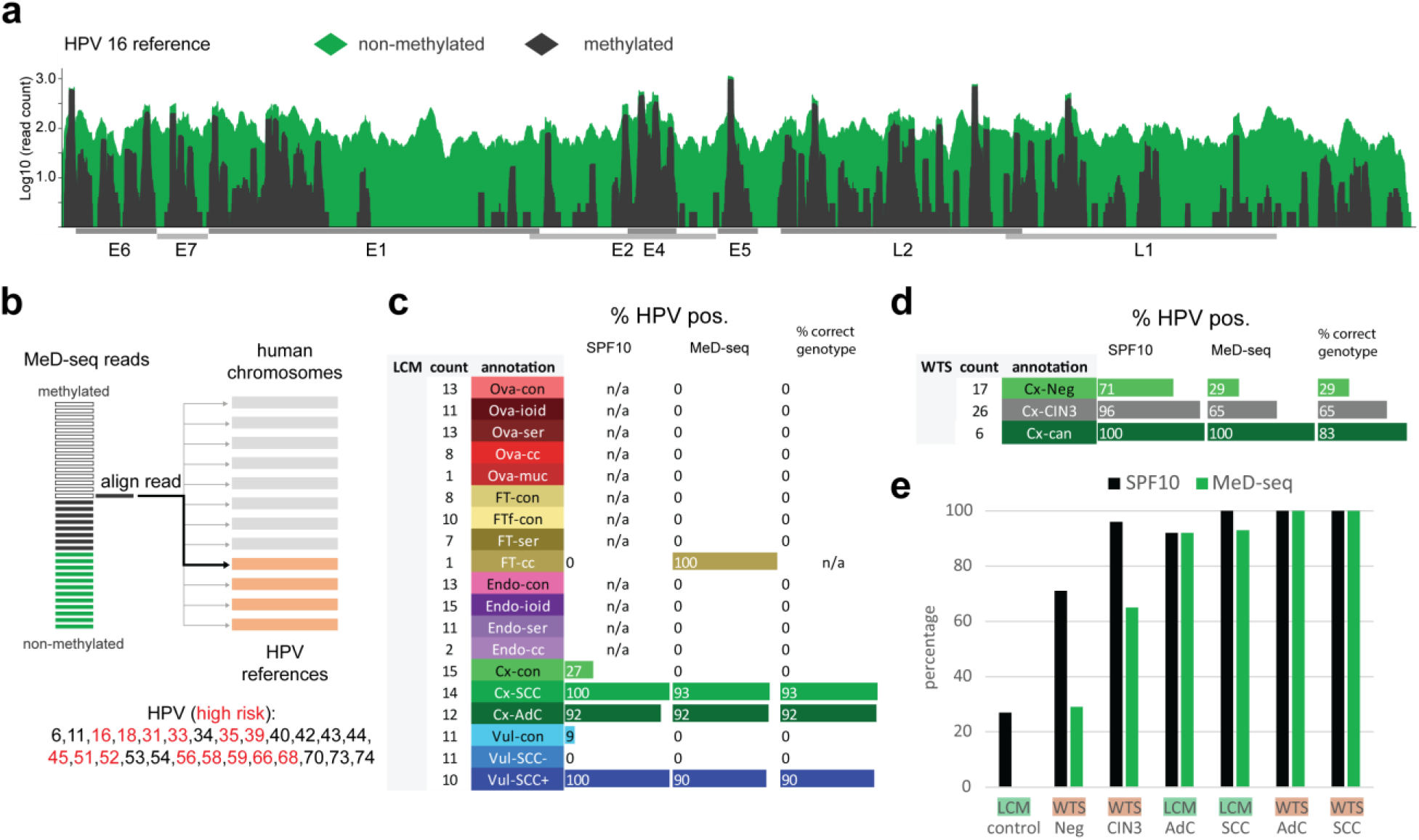
HPV genotype detection using MeD-seq. (a) MeD-seq read coverage of all methylated (black) and non-methylated (green) reads, from all samples in (3c), stacked on the HPV 16 genome. (b) Viral DNA detection pipeline. Methylated and non-methylated MeD-seq reads were used to map against the human genome and various HPV genomes at the same time. (c) Percentage of HPV genotypes detected by SPF10 and MeD-seq. Percentage of corresponding detection of same HPV genotype using MeD-seq and SPF10 is shown (right). (d) As in (3c) but now for cervical negative (Cx-Neg), cervical CIN3 lesion (Cx-CIN3) and cervical cancer (Cx-can) WTS. (e) Bar plot comparison showing overview of results presented in (c) and (d).

To test whether HPV could also be detected in whole tissue slices (WTS) we applied SPF10 and MeD-seq analysis on WTS of cervix SCCs, cervix AdCs, CIN3 cervix cancer pre-stage lesions and controls. This analysis revealed a high percentage of controls and CIN3 samples with high-risk HPV, and a 100% detection of high-risk HPV in cervix SCC and AdC WTS samples applying the SPF10 test (Fig. 3c-d). This was different when we applied MeD-seq analysis where we found an increasing number of positive samples from pre-stage to full cancer, which can be attributed to the fact that the SPF10 test detects integrated and non-integrated HPV, whereas MeD-seq only detects the methylated HPV genome that is likely only present after integration of HPV into the host genome.

### Defining cancer associated DNA methylation marker regions

To define DNA methylation marker regions that best predict cancer-associated changes we applied a genome-wide sliding window approach and defined different types of DMRs; general cancer DMR’s that are methylated in all cancers and not in healthy control tissue, regional cancer DMR’s that are methylated in a subset of cancers and not in other cancers and healthy control tissue, and specific cancer DMR’s that are only methylated in one specific cancer type and not in other cancers and healthy tissues. In addition, we identified DMR’s that were hypermethylated in all controls or specific healthy controls. We focused on small 80 bp regions to facilitate development of PCR-based assays on selected DMRs. For each individual DMR a threshold was calculated, and samples were scored in a binary fashion (0 below and 1 above the threshold).

Individual DMRs were ranked based on an area under the curve (AUC) score and the 50 highest ranked DMRs were selected per comparison (Fig. 4a). Using all selected DMRs, we performed hierarchical clustering analysis with all individual samples (Fig. 4b, Supp. Fig. 1c). In addition, we generated a UMAP based on all defined specific DMRs (Fig. 4c). These analyses revealed that all cancers could be discriminated from controls, but also indicated that the power to distinguish cancer groups and/or subtypes varied between subtypes and regions of origin. Cancers originating from the vulva or cervix could clearly be distinguished from endometrium, fallopian tube or ovarian cancers. In addition, DMRs could be identified which distinguish most specific cancer subtypes and control tissues (Fig. 4b,c). In contrast, several cancers originating from the fallopian tube (serous) or ovary (serous and endometrioid) could not be separated. Examination of individual DMRs confirms the clear distinction in methylation levels between cancers, and healthy tissues, and different cancer types, providing a powerful resource for assay development (Fig. 4d).

**Figure 4.**
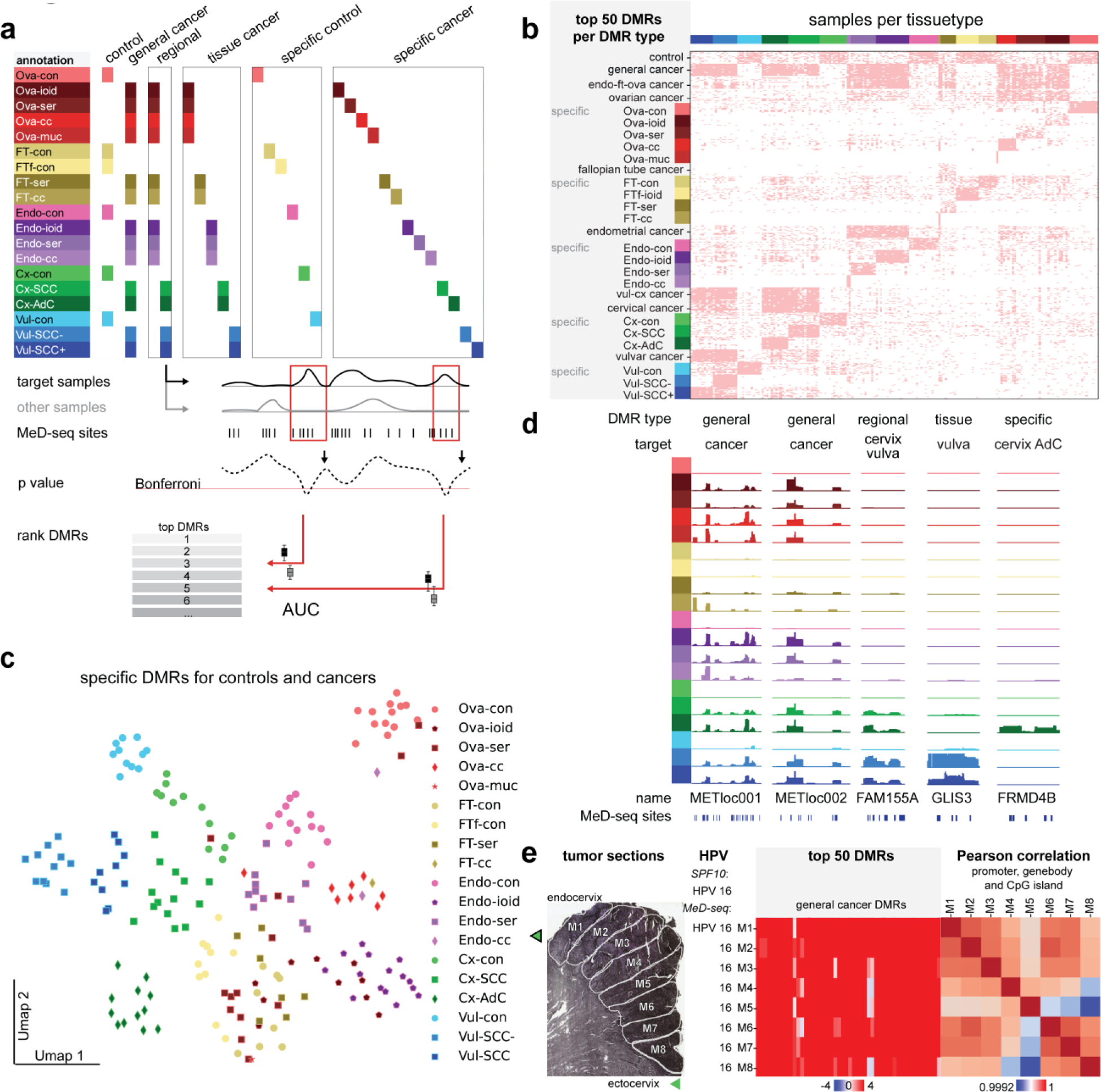
Cancer associated DNA methylation marker regions. (a) DMR detection workflow. Predefined DMR types, with a target sample group vs all other samples defining; general control DMRs, general cancer DMRs, regional DMRs, tissue cancer DMRs, specific control DMRs, specific cancer type DMRs. Using a sliding window a DMR is selected when the p-value of the t-test between the target and the other samples is below the Bonferroni p-value. The resulting DMRs are then ranked on the area under the curve (AUC) score. (b) Heatmap of the top 50 DMRs per comparison (DMR types), using a DMR threshold on MeD-seq read counts per DMR, values are either 1 (pos/red) or 0 (neg/white). (c) UMAP generated with all samples using the top 50 specific control and cancer DMRs. (d) Average MeD-seq coverage for two general cancer DMRs (METloc001/METloc002), one regional DMR (FAM155A), one tissue cancer DMR (GLIS3) and one Cx-AdC specific DMR (FRMD4B). (e) Eight LCM tumor sections with anatomical relation to endo- and ectocervix (left panel). MeD-seq HPV genotype detection per tumor section (middle left panel). Z-score heatmap of the top 50 general cancer markers in different sections (middle right panel), compared to cervix control samples. Pearson correlation of MeD-seq results (right panel) obtained with different tumor sections within the tumor, using promoter, gene body and CpG-island DNA methylation.

We next wanted to test whether the ability of our defined DMRs to detect cancer was independent of cancer heterogeneity. We therefore used LCM to isolate tumor DNA from eight different zones in two different SCCs, and subsequently performed MeD-seq on each zone (Fig. 4e, Supp. Fig. 1d). Genome-wide DNA methylation analysis involving promoter, gene body and CpG-island methylation uncovered modest differences between the different zones, and Pearson’s correlation analysis indicated that changes in DNA methylation to be most prominent in the central region of both tumors (Fig. 4e, Sup. Fig. 1d). In contrast, the 50 DMRs marking all cancers, defined above, displayed consistent methylation in all zones. Similarly, HPV MeD-seq read-count analysis revealed high-risk HPV subtype present in nearly all zones, indicating that the most predictive changes in DNA methylation, as well as the presence of integrated HPV, are stable changes associated with tumorigenesis.

### Development and validation of targeted assays

To enable straightforward cancer detection that can be easily implemented in a clinical and/or population screening setting, we developed a qMSP assay capable of detecting all gynecological cancers. We selected the most promising DMRs with the highest discriminating power, by selecting the DMRs with the highest area under the curve (AUC) values based on our MeD-seq data. For our DMR selection we compared all five (vulva, cervical, endometrial, fallopian tube and ovarian) types of cancer to controls, or a limited dataset consisting of only cervical and endometrial cancers and controls. From the full dataset comparison, we selected two intergenic regions (chr1:145475713-145476399 and chr3:147384749-147385097) that, due to the absence of a gene annotation were labeled as “METloc001” and “METloc002” respectively. Two intragenic regions (inside ARID3C; chr9:34623552-34624056 and ARL5C; chr17:39165088-39165444) were selected from the cervical/endometrial-only dataset. To estimate the potential discriminatory power of the four qMSP assays, we first analyzed MeD-seq data from regions targeted for qMSP. Four qMSP assays were developed and tested on WTS sections of tumor and healthy tissues including both samples that we used to generate the MeD-seq database as well as novel samples. Our results showed that both METloc001 and METloc002 can distinguish cancer from healthy tissues, and showed an increasing trend of DNA methylation levels in cervical pre-lesions (CIN1 to 3) (Fig.5a). The ARID3C and ARL5C qMSP assays show results similar to those found using METloc001 and METloc002 for cervical and endometrial cancer, however for detection of vulva, fallopian tube and ovarian cancer, ARID3C and ARL5C display less consistent results. For instance, ARID3C can’t properly distinguish fallopian tube cancers from healthy fallopian tube controls or vulva cancers from controls. When comparing AUC values and test accuracy, METloc001 and METloc002 outperform ARID3C and ARL5C (Fig. 5b). Combinatorial analysis using two marker regions showed that the METloc001/METloc002 combination is most powerful with a sensitivity of 98% and a specificity of 99% on WTS samples when detecting all gynecological cancers that we tested (Fig 5b/Table2). This clearly shows the importance of taking all types of gynecological cancers and healthy tissue samples into account when selecting a DNA methylation marker region.

**Figure 5.**
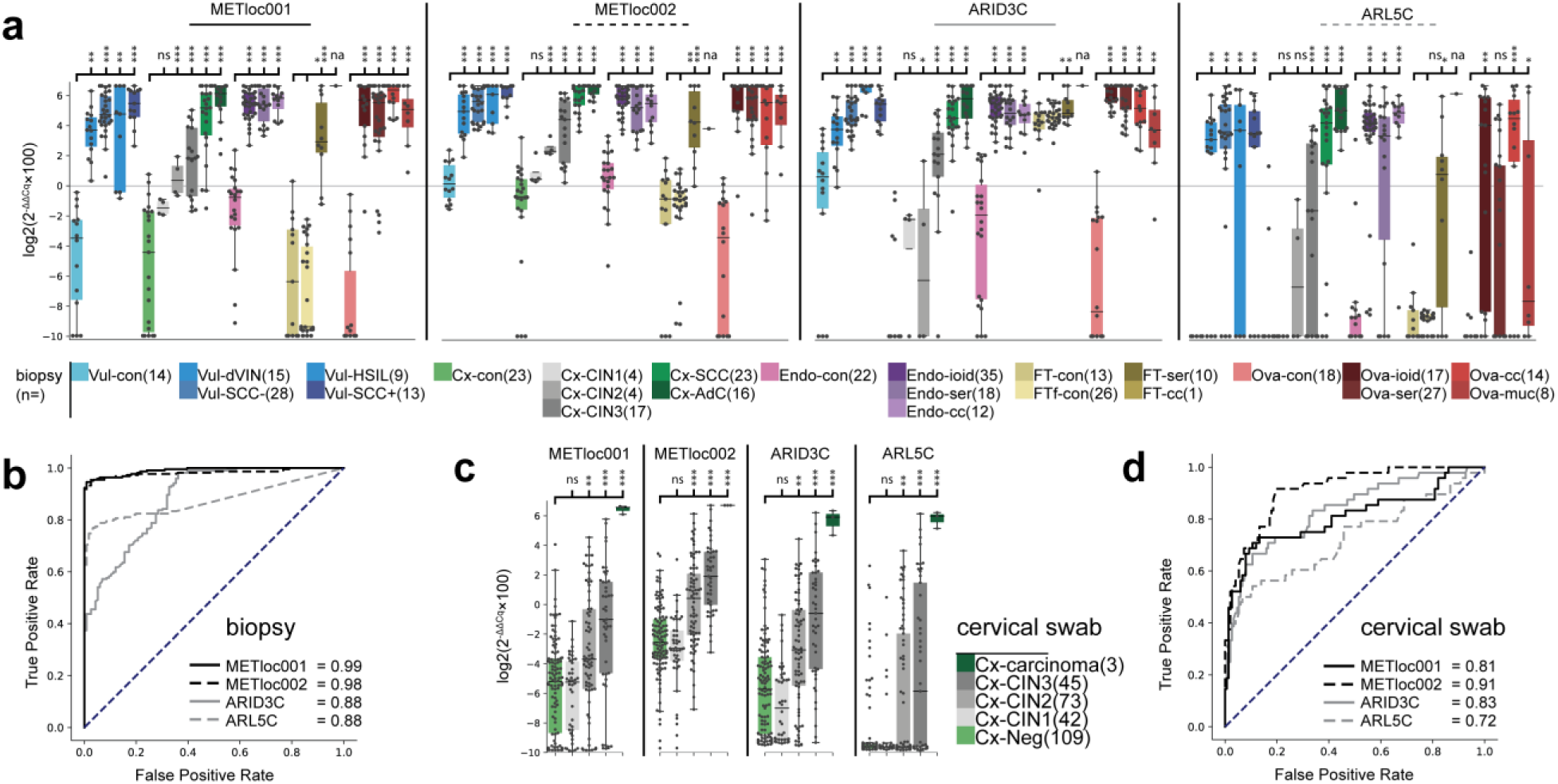
Development and validation of targeted assays for cervical screening. * p < 0.05, ** p < 0.01, *** p < 0.001 (a) qMSP results of four general cancer biomarkers on biopsy or resection samples. (b) ROC curve with AUC values per biomarker on biopsy samples. (c) qMSP boxplot results per biomarker on cervical swab samples from the EVAH study. (d) ROC curve with AUC values per biomarker on cervical swab samples.

Next, we tested all four qMSP assays on cervical swab material from the EVAH study, a multicenter study on a cytology-screened referral population [7]. Similarly to the WTS results, all four assays can clearly distinguish cancers from controls on cervical swabs. In addition, qMSP analysis of cervical swabs also revealed an increase in DNA methylation levels for all four marker regions, when comparing CIN1 to CIN3 cervical pre-cancer lesions (Fig.5c). The EVAH study aimed to detect cancer and CIN3 lesions. Therefore, we determined a CIN3+ threshold against ≤CIN1. Despite the fact that selection of all four marker regions was based on MeD-seq data that included cervical cancers, the METloc001/METloc002 combination also outperformed other combinations with a sensitivity of 94% and a specificity of 79% for detection of CIN3+ lesions suggesting that the addition of other gynecological cancers in the selection process improved cervical cancer detection performance (Fig. 5d/Table 2). A previous study involving the EVAH cohort applied a qMSP assay, whereby a CIN3+ threshold was determined by setting the specificity at 70% (which resulted in a sensitivity of 77,8 %), applying this strategy on the METloc001/METloc002 combination increased sensitivity from 94% to 96% [7].

Because CIN lesions consist of a heterogeneous group of lesions, that can either regress, persist or progress to cancer, we set out to look at the consistency of our four markers in the EVAH study (Supp. Fig. 2a). Consistency in qMSP results (all four assays positive or all four negative) was found for the vast majority of control (83,3%), CIN1 (88,1%) and cancer (100%) samples. In addition, CIN2/3 lesions also display consistent results for the majority of samples, 75,4% of CIN2 lesions and 57,8% of CIN3 lesions (Fig. 6a,b).

**Figure 6.**
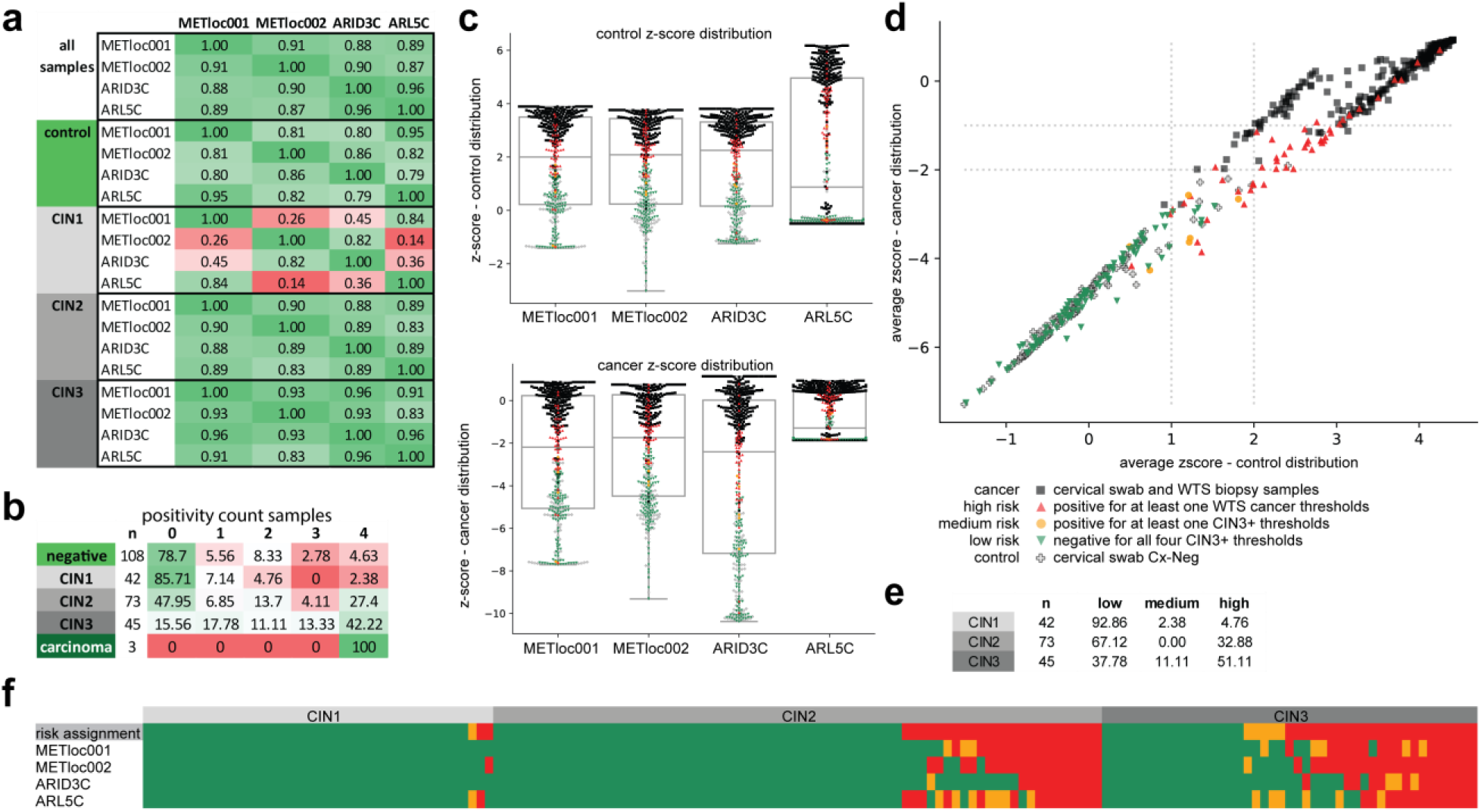
Risk assessment of CIN lesions. (a) Pearson correlation on EVAH swab samples, comparison between four biomarkers. Correlation matrix for all samples, negative control samples and per CIN lesion. (b) Percentage of samples per positivity count for every swab subtype. (c) Boxplots per biomarker, y-axis z-score based on μ and σ of swab and biopsy control (top) or cancer (bottom) samples. CIN lesions subdivided into low, medium and high risk (see legend d). (d) Scatterplot of EVAH and biopsy cancer samples. x-axis z-score calculated using the mean and standard deviation of negative control samples. y-axis z-score calculated using μ and σ of cervical swab samples and biopsy or resection cancer samples. CIN lesions are subdivided into low, medium and high risk. Gray dotted guidelines for z-score of [-2, -1, 1, 2]. (e) Percentage of cervical swab samples categorized in low, medium and high risk; per CIN lesion type. (f) Risk assignment and biomarker risk result comparison, samples are colored green/low risk, orange/medium risk or red/high risk.

To perform risk stratification on controls, we calculated Z-score values of all four qMSP assay results for CIN1-3 and cancer samples. These Z-scores were either calculated using mean and standard deviation from control samples (X-axis) or mean and standard deviation from cancer samples (Y-axis) (Fig. 6c,d). Control samples were cervical swab material from the EVAH study, whereas cancer samples were cervical swabs from the EVAH study (n=3) supplemented with WTS of gynecological cancers (n=222).

Afterwards, two different thresholds were calculated using ROC curves, a cancer detection threshold and a CIN3+ detection threshold. Samples were then classified as either high risk (above cancer threshold for at least one marker), medium risk (above CIN3+ threshold for at least one marker) or low risk (below CIN3+ threshold for all four markers) (Fig. 6e,f). This resulted in >90% of CIN1 lesions being low risk, and >50% of CIN3 lesions being high risk. Using these medium+-risk classification thresholds and focusing solely on medium risk CIN lesions instead of all CIN3 lesions, we could increase the performance of the individual markers and increase the specificity of METloc001/METloc002 combination from 79% to 93%, without sacrificing test sensitivity (94% to 93%) (Table 2, Supp. Fig. 2b).

### Biomarkers for cancer subtypes and non-gynecological cancers

Besides general biomarkers detecting all gynecological cancers, we selected three DMR regions based on MeD-seq data, which detect a subset of gynecological cancers, for qMSP assay development. An intergenic region named METloc009 (chr10:23173346-23173626) was selected as a biomarker for vulva and cervical cancers only, and genomic regions in GLIS3 and FRMD4B were selected as specific biomarkers for either vulva cancer or cervical adenocarcinoma respectively. METloc009 qMSP analysis on biopsy material revealed that vulvar and cervical cancers could be clearly distinguished from control samples (Fig. 7a-b). Although GLIS3 and FRMD4B can detect vulvar cancer and cervical adenocarcinoma respectively, differences in methylation between cancer and associated healthy tissues are less pronounced than what we observed when using METloc009. In addition, GLIS3 which was selected as a biomarker for vulva cancer, also shows elevated DNA methylation levels in cervical squamous cell carcinomas, consistent with both cancers being squamous cell carcinomas (Fig. 7a).

**Figure 7.**
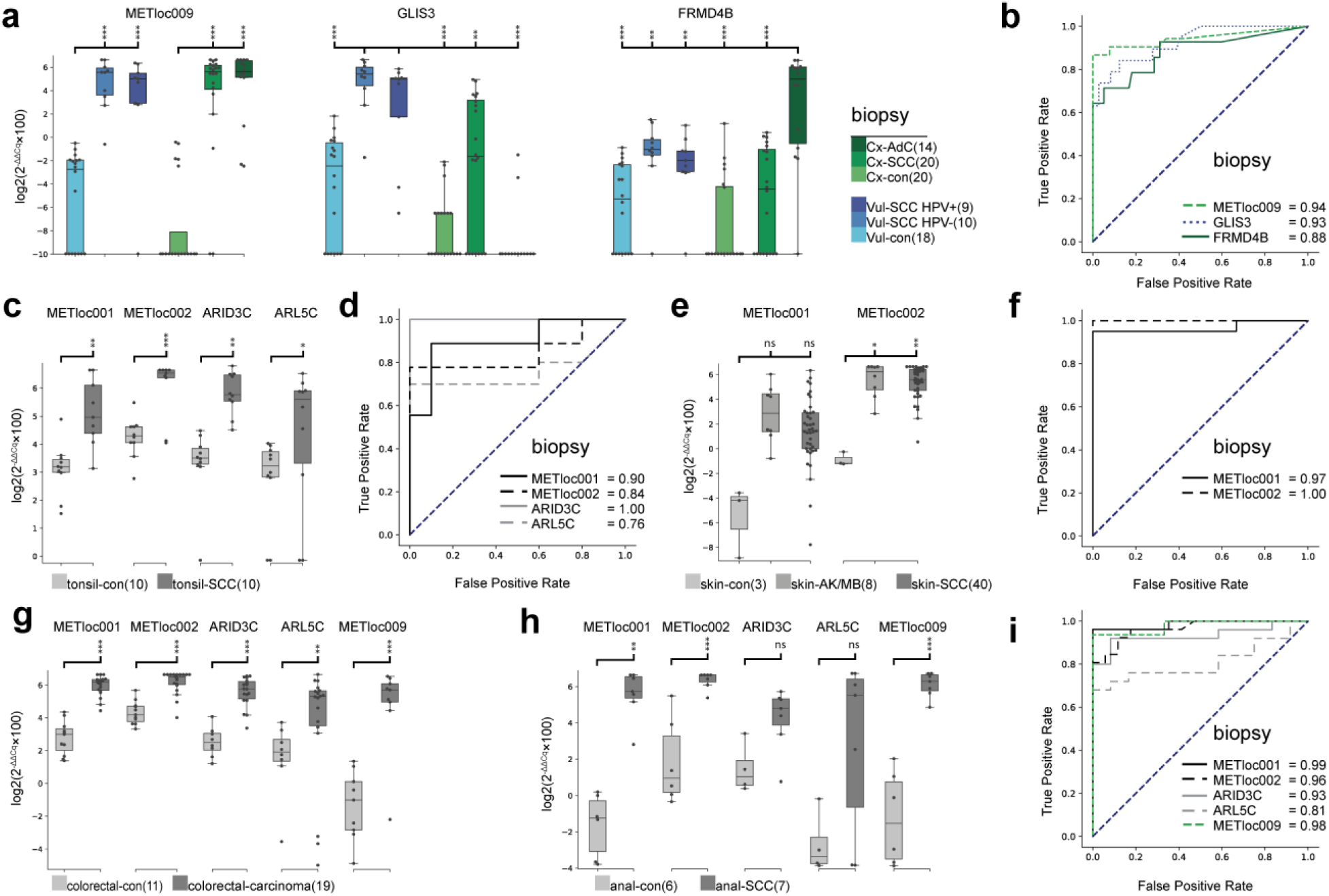
Biomarkers for cancer subtypes and non-gynecological cancers. * p < 0.05, ** p < 0.01, *** p < 0.001 (a) qMSP results for 3 biomarkers tested on biopsy samples. METloc009 detecting vulvar and cervical cancer (regional DMR); GLIS3 detecting vulvar cancer (tissue cancer DMR) and FRMD4B cervical AdC (specific cancer DMR). (b) ROC curve with AUC values per biomarker on vulvar and cervical biopsy samples. (c) qMSP results of four general cancer biomarkers on biopsy tonsil samples. (d) ROC curve with AUC values per biomarker on biopsy tonsil samples. (e) qMSP results of two general cancer biomarkers on biopsy skin samples. (f) ROC curve with AUC values per biomarker on biopsy skin samples. (g) qMSP results of four general (METloc001, METloc002, ARID3C, ARL5C) and one regional (METloc009) cancer biomarkers on biopsy colorectal samples. (h) qMSP results of biomarkers tested in (g) on biopsy anal samples. (i) ROC curve with AUC values per biomarker on biopsy colorectal and anal samples.

As several of our biomarkers detect multiple cancers, we wanted to test whether METloc001, METloc002, ARID3C, ARL5C and METloc009 could also be applied to detect a diverse set of non-gynecological cancers. METloc001 and METloc002 were used to detect tonsil, skin, colorectal and anal cancer, ARID3C and ARL5C were used to detect tonsil, colorectal and anal cancer, and finally METloc009 was used to detect colorectal and anal cancer. All cancer DNA was isolated from biopsy material. Our results indicate that, notwithstanding the selection of all biomarker regions from a set consisting only of gynecological cancers, METloc001, METloc002, ARID3C and ARL5C could easily distinguish tonsil, colorectal, and anal cancer from associated healthy tissues (Fig. 7c,d,g-i). In addition, METloc001 and METloc002 could distinguish actinic keratosis (AK), Morbus Bowen (MB) lesions, and skin cancer from associated healthy controls (Fig. 7e,f). Even METloc009, selected to detect vulva and cervical cancer could distinguish colorectal and anal cancer from associated controls (Fig. 7g-i). These findings indicate that our DNA methylation markers aimed at detection of gynecological cancers also detect unrelated cancers in other regions with another cell type of origin, likely reflecting epigenetic changes affecting common pathways in cancer development.

## Discussion

In this study we applied MeD-seq to obtain genome-wide DNA methylation profiles of gynecological cancers, which we used as a foundation to develop several novel cancer markers, including two promising markers for the detection of all gynecological cancers. We identified differentially methylated regions that are cancer-type specific, specific for sub-groups of cancers or common to all gynecological cancers. We selected the most predictive regions that are differentially methylated in all gynecological cancers for development of a qMSP assay, and applied this assay to detect cancer using biopsies or swabs of women with cervical cancer and different cervical pre-cancer stages. Our studies reveal a high sensitivity and specificity for detecting CIN3+ lesions in cervical biopsies and swabs. Cervical swabs from the EVAH study tested with METloc001/ METloc002 showed a sensitivity of 94% and specificity of 79% for CIN3+ lesions, an improvement compared to another qMSP assay performed on the EVAH study (sensitivity 77,8%, specificity 69,3%) [7]. In addition, we found that the specificity could be further increased by focusing on medium+-risk CIN lesions.

We found that markers which were developed using the set of all gynecological cancers outperformed those that had been selected from among a subset of cancers.

Surprisingly, testing of our general markers METloc001 and METloc002 on other cancer types that were not part of our current MeD-seq database showed that these general markers could still distinguish cancer from healthy tissues, despite these cancers not being in the original discovery set. However, we expect that a genome wide DNA methylation approach using MeD-seq on tumor material from these non-gynecological cancers and their associated healthy tissues will yield novel biomarkers that are able to distinguish cancer from healthy tissues more efficiently.

Close inspection of METloc001 and METloc002 locations using the UCSC database indicates both regions to be enriched for CG dinucleotides, and overlap with regulatory regions (Remap database) that show long range genomic interactions in different cell types (Genehancer). These findings suggest that METloc001 and METloc002 are likely enhancer elements that are very frequently methylated in a wide range of cancers.

Similar enrichment results regarding CG dinucleotides and regulatory regions can be found when inspecting the intragenic marker regions in ARID3C and ARL5C. We looked for binding sites in the four aforementioned regions and found a plethora of regulatory factors. Whereas METloc001 is only enriched for TATA-binding protein (TBP) and BRF1 binding sites (both associated with control of gene transcription [16, 17]), the METloc002, ARID3C and ARL5C regions are enriched for a wide range of regulatory factors. These factors include CTCF and RAD21 for METloc002 and ARL5C, which are known to be involved in genome organization and transcription regulation [18, 19], whereas FOXA1 and BRD4 are enriched at ARID3C and are known to regulate cell differentiation [20, 21]. Despite our general markers being associated with different biological pathways, they are methylated in a wide range of cancer types and are also consistently methylated in different parts of the same tumor. This suggest that although different driver mutations in a range of cancer types might initiate carcinogenesis, our cancer biomarkers are downstream of these genomic events and are generally present in cancers. These results show the potential power of our biomarker assays but also of DNA methylation in general for detection of cancer and its precursor lesions.

The main four biomarkers presented in this study (METloc001, METloc002, ARID3C and ARL5C) show high consistency in hypermethylation or hypomethylation, when qMSP assays are performed on cervical biopsies and cervical swabs. This was to be expected regarding tumor material, since all four biomarkers were selected on biopsy material in our database. Nevertheless, we were surprised to see that this consistency extended also to the CIN1-3 precursor lesions. CIN lesions can regress, including the more advanced CIN3 lesions, but predicting which CIN lesions regress, persist or progress towards cancers is currently not yet possible. Our results show that the majority of CIN1-3 lesions are either positive or negative for all four of our qMSP biomarker assays. This inter-marker concordance could potentially predict future outcomes of CIN lesions. To test this hypothesis, qMSP assays targeting METloc001, METloc002, ARID3C and ARL5C should be performed on cervical swab and/or biopsy material from CIN lesions with patient follow-up information.

The present qMSP assays used for the detection of cervical cancer are targeted towards detecting CIN3+ or CIN2+ lesions in clinical studies. Biomarker qMSP assay thresholds focusing on cancer detection generally display a much higher sensitivity and specificity. The common aim to also detect all CIN3 or even all CIN2 in most studies leads to a dramatic decrease in assay specificity when used in screening, due to a large number of CIN2-3 lesions having DNA methylation levels similar to healthy tissue samples. Therefore, we constructed a novel risk classification that aims at detecting medium to high-risk CIN lesions, regardless whether they are CIN1, CIN2 or CIN3 based on our four biomarkers showing strong inter-marker consistency. This CIN lesion risk stratification leads to a full molecular classification system of cervical swab samples into low, medium and high-risk groups. In addition, we adjusted the thresholds for each individual qMSP marker assay to detect medium and high-risk lesions. This led to a strong increase in specificity (79% to 87%) while retaining high sensitivity (94% to 93%) when focusing on medium-high risk detection instead of CIN3+ detection using our biomarker qMSP assay consisting of METloc001 and METloc002. If this approach can be validated using patient follow-up information as described above, a CIN1 lesion which is classified as “methylation high-risk” could be considered of higher risk than a “methylation low risk” CIN3 lesion. This could pave the road to a DNA methylation-based methodology of detecting women at risk in molecular pathology.

MeD-seq is able to detect DNA fragments originating from the HPV genome, which was confirmed by the SPF10 assay. In addition, classification of the correct HPV subtype was very consistent between SPF10 and MeD-seq results. However, a large discrepancy between SPF10 and MeD-seq was found in healthy cervical control tissues, where the HPV detection rate by MeD-seq is much lower than SPF10 (29% versus 71% respectively). We speculate that HPV detection by MeD-seq might be dependent on HPV integration into the host genome, where the integrated HPV becomes chromatinized and (further) methylated.

Several methods are available that provide genome wide profiling of DNA methylation, however the ability of MeD-seq to enrich for and sequence the methylated regions of the genome leads to a strong reduction in sequencing costs. Because MeD-seq requires a relatively low amount of DNA as input (making it compatible with LCM) we could generate a large DNA methylation database of cancers and control material from the female reproductive tract, with high target cell enrichment at low cost. Since MeD-seq is compatible and easily applied to liquid biopsies [22], an interesting application of the current MeD-seq database is to detect and classify gynecological cancers in cell free DNA from blood plasma. Extending the MeD-seq database could further enable biomarker discovery to detect a much wider range of cancer types with custom qMSP assays, while also creating a DNA methylation database for the direct detection, classification and monitoring of disease status and treatment success (MRD) using MeD-Seq.

## Materials and methods

### Cervical swabs

The EVAH study is a prospective multicenter observational cohort study 16 conducted in Voorburg, The Netherlands, and Barcelona, Spain. Inclusion criteria were (i) an abnormal cyto-logical test result (≥ASC-US) and (ii) 18 years of age or older. The criteria of exclusion were (i) previous diagnosis of ICC, (ii) history of surgery to the cervix or previous pelvic radiotherapy, (iii) current pregnancy or pregnancy in the previous 3 months, (iv) current breastfeeding or breastfeeding in the previous 3 months and (v) insufficient material for hrHPV testing and methylation analysis.

A trained physician collected a cervical smear using a Cervex-Brush (Rovers Medical Devices B.V., Oss, The Netherlands). The Cervex-Brush was rinsed in 20 mL of ThinPrep medium (Hologic, Marlborough, MA). Remnant of the original sample was spun down and the cell pellet was re-suspended in 2 ml of ThinPrep and this was stored at 4 ֯C until further testing.

0.5 ml of the stored sample was spun down and the cell pellet was re-suspended in 0.5 ml of 50 mM Tris-HCl pH 8.0. DNA was extracted with the NucliSENS® easyMAG® (bioMérieux) using the ‘Generic Protocol 2.0.1’ (input volume 500 µl and elution volume 50 µl) according to the manufacturer’s instructions. After the extraction of genomic DNA, the concentrations and quality of the extracted DNA were determined in the DNA QC qPCR.

### Biopsy samples

Coded patient samples were obtained from a clinical study, which was approved by the Medical Ethics Committee from the Erasmus University Medical Center (MEC-2020-0846). All biopsy blocks were sectioned according to the sandwich cutting procedure: a 4 μm section for diagnosis (H&E before); 16 μm sections for LCM-MeD-seq analysis; a set of 3-6 × 8 μm (24 μm) sections for WTS-qMSP analysis, and finally, a 4 μm section for pathological confirmation (H&E after).

### Laser capture microdissection (LCM) for target cell enrichment

For good-quality genome-wide DNA methylation input of a purest possible population of target cells is required. LCM is an accurate method for isolating these specific target cells out of complex heterogeneous tissue [23]. To improve tissue adhesion to the membrane, PALM membrane slides 1.0 PEN were prior to use UV treated (254 nm) and coated with poly-L-lysine (0.1%) according to manufacturer’s protocol. 16 µm sections of formalin-fixed paraffin-embedded (FFPE) tissues were cut, mounted on the UV and poly-L-lysine pretreated membrane slides and incubated at 60°C for 2 hours. Next, the sections were dewaxed, dehydrated and stained with haematoxylin. After digital imaging of the stained slides, regions of interest were annotated by a pathologist. The annotated regions must contain the highest possible percentage of desired target cells (preferably >70%) surrounded by only normal tissue cells (and preferably no other pathological structures). Using LCM, the percentage of desired cells in the region of interest (precancer lesion, cancer lesion or normal epithelium) can be increased and regions with unwanted cell types/cell structures can be removed. The annotated regions were cut by a laser beam using the Leica Laser Microdissection system and collected in tubes. For the MeD-seq analysis we need at least 10 (preferably 50) ng DNA per 8.66 μl of a sample. For this we dissected one or more regions that together were 5-20 mm2.

### DNA extraction FFPE tissue

DNA was extracted using our Proteinase K protocol. The first steps differ between WTS and LCM samples. To each WTS sample 700 μl of mineral oil was added. The samples were vortexed and spun followed by an incubation for 3 min at 70°C and 5 min at room temperature. Then 100 μl of 1 mg/ml Proteinase K was added. To each LCM sample 50 μl of 1 mg/ml Proteinase K and 200 μl of mineral oil were added. Thereafter, all sample types were incubated 1-2 h at 70°C. The samples were then gently mixed and incubated overnight for 16 h at 70°C. If the tissue was not completely dissolved, 5 µl 10 mg/ml Proteinase K was added and the samples were incubated for 6 h at 70°C. Afterwards, Proteinase K was inactivated by 10 min at 95°C and the mineral oil was removed. Samples were stored at -20°C. After the extraction of genomic, the concentrations and quality of the extracted DNA were determined using the DNA QC qPCR.

### DNA QC qPCR

The quantity and quality of amplifiable human DNA isolates was determined using a multiplex inhouse qPCR, the DNA QC qPCR, amplifying two fragments of different size (human single copy genes RNase P-68bp and ACTB-145bp) and including an internal control (IC) to detect for PCR-inhibition. A standard curve consisting of serial dilutions with known RNase P concentrations was used to calculate the DNA concentration of each sample. The difference between the Cq-values of the two fragments (ΔCq = Cq 145 bp ACTB fragment - Cq 68 bp RNaseP fragment) provides an effective quality control for identifying degree of fragmentation. The qPCR mix contained a plasmid spiked at a fixed concentration that functioned as an internal control to detect PCR-inhibition. An elevated Cq value of the IC target is an indication of the presence of PCR inhibition.

The DNA QC was performed in a final reaction volume of 12.5 µl, containing designed primers/probe sets, PhHV plasmid and the extracted DNA in a PCR Master Mix, and under the following conditions: 95.0°C for 5 minutes followed by 45 cycles of 95.0°C for 15 seconds and 60.0°C for 50 seconds. Fluorescence data was collected at the end of each annealing/extension step for determination of the Cq values. The multiplex was performed on the CFX96 7 Real-Time PCR System. Cq values were measured at fixed thresholds for fluorescence.

### SPF10

Ten µl of the undiluted extracted DNA or the lowest dilution without PCR inhibition was used for the detection of HPV DNA by PCR amplification using the HPV SPF10 PCR/LiPA25 (version 1) system (Labo Biomedical products, Rijswijk, The Netherlands)[24, 25]. The SPF10 PCR primer set amplifies a small fragment (65 base pair) from the L1 open reading-frame of the HPV genome.

The DNA enzyme immunoassay (DEIA) test was used to detect amplified HPV DNA products using a cocktail of universal probes detecting DNA from at least 69 HPV genotypes [26]. DEIA test reactions were measured by optical densities at 450 nm (OD450). The interpretation of the DEIA test result was done by comparing the OD450 of the HPV PCR products to the OD450 value of a DEIA borderline control sample (DEIA negative if OD450<0.160, DEIA borderline if OD450 between or equal to 0.160 and OD450 DEIA borderline control, and DEIA positive if OD450> OD450 DEIA borderline control).

HPV genotyping was performed on all SPF10 PCR DEIA-borderline and DEIA-positive samples using reverse hybridization line probe assay membrane strips (LiPA25) in a ProfiBlot 48 analyser (Tecan Austria GmbH, Salzburg, Austria) to identify 25 anogenital HPV genotypes: HPV6, 11, 16, 18, 31, 33, 34, 35, 39, 40, 42, 43, 44, 45, 51, 52, 53, 54, 56, 58, 59, 66, (68 or 73), 70, and 74. Positive LiPA25 results were visualized by a color substrate reaction immobilized on the probe lines and the final SPF10 results were interpreted as follows: 1) positive for a specific HPV genotype, 2) positive for HPV by DEIA but un-typable genotype by LiPA25 or 3) no HPV detected.

### MeD-seq assay

MeD-seq assays were performed as previously described [14]. In short, genomic DNA and plasma-derived cfDNA were digested with the methylation-dependent restriction enzyme LpnPI (New England Biolabs, Ipswich, MA) generating 32 bp DNA fragments containing the methylated CpG in the middle. All samples were sequenced on the Illumina NextSeq2000 platform.

### MeD-seq data analysis

Custom python scripts were used to process acquired DNA methylation profiles using the following python version and packages: python v3.8.5, numpy v1.19.1, scikit-learn v0.23.2, scipy v1.5.2, matplotlib v3.3.1, seaborn v0.11.0, umap-learn v0.5.1. Raw fastq files were subjected to Illumina adaptor trimming and reads were filtered based on LpnPI restriction site occurrence between 13–17 bp from either the 5′ or 3′ end of the read. Reads that passed the filter were mapped to hg38 using bowtie2 version 2.3.3 and BW files were generated using wigToBigWig version 4 for visualization in IGV. Genome-wide individual LpnPI site scores were used to generate read count scores for the following annotated regions: transcription start sites (TSS, 1 kb before and 1 kb after), CpG-islands and gene bodies (1 kb after TSS till TES). Gene and CpG-island annotations were downloaded from ENSEMBL (Homo_sapiens_hg38.GRCh38.79.gtf, www.ensembl.org).

Differentially methylated regions (DMRs) were detected using custom Python scripts. In addition, a sliding window technique was used to detect DMRs and the Chi-squared test on read counts was used for statistical testing with a Bonferroni correction. A Bonferroni corrected p-value ≤0.05 was considered statistically significant. Z-score transformation of the read count data was applied for unsupervised hierarchical clustering analyses. DMRs were identified between methylation profile from cervical control samples compared to methylation profiles of WTS, WTS<=70% and LCM samples.

For genome wide comparison we used promoters(TSS), gene bodies and CpG-island as genomic locations. These genomic locations were filtered, regions locations were used if at least three samples had zero reads, at least three samples had reads and any sample can’t have a maximum of 10 reads. A umap was generated based on promoters(n=19185), gene bodies(n=3476) and CpG-islands(n=14078). These same locations were used to compute the Pearson correlation between samples.

All boxplots are visualized using packages matplotlib and/or seaborn. Significance was calculated with an independent ttest implemented in scipy. Pearson correlation and z-score was computed using scipy package.

To perform multiple analysis at ones we define DMR types, DMR types are groups of sample types that are analyzed as one target sample group versus the other remaining samples. DMR types were constructed based on biological assumptions or logic association, based on being a cancer or control, regional anatomy, tissue collection or unique samples type.

Using a sliding window of 80 base pairs for every MeD-seq site per DMR type a p-value is calculated using Mann–Whitney U test (scipy package), if the p value is above the Bonferroni threshold (0.1/MeD-seq sites count) it was discarded. Resulting MeD-seq site windows (DMRs) per DMR-type were ranked using AUC score.

To determine the presence or absence of methylation profiles associated specifically with all cancers or unique cancer for example, receiver operating characteristic (ROC) curves were calculated for each sample for each individual DMR. ROC curves of target samples compared to other remaining samples, were used to calculate the optimal threshold (using the “scikit-learn” package Python) for each individual DMR. Samples above the threshold scored ‘1’, samples under the threshold scored ‘0’ (binary score).

Heatmaps were generated using the binary scores for all the different DMR types.

For HPV genotype detection using MeD-seq we align the reads using bowtie2 against a combined reference file. The combined reference file contains all the human hg38 reference chromosomes and genotype references of HPV that can be detected in the SPF10 test. The generated alignment file by bowtie2 is then scanned for reads that were aligned to a HPV reference, these reads are counted for every HPV genotype and scored methylated or non-methylated. A minimal threshold of three reads was applied to detect the HPV genotype. If SPF10 had a un-typable results it was considered as a negative result, because the MeD-seq analysis can only detect HPV genotypes that have their reference in the combined reference file.

### Bisulfite Conversion

To perform the quantitative methylation-specific PCR, extracted DNA was bisulfite-converted using the EZ DNA Methylation kit (Zymo Research, Orange, CA, USA) according to the manufacturer’s instructions, converting unmethylated cytosines into uracils while methylated cytosines remain unchanged. Elution was performed in 12.5 µl with final concentration of 10 ng/µl for biopsy samples and 20 ng/µl for cytology samples.

### Quantitative methylation-specific PCR (qMSP)

Bisulfite converted samples were analyzed for methylation using the multiplex qMSP assay, which targets selected genes and the reference gene β-actin (ACTB). The qMSP was performed in a final reaction volume of 12.5 µl, containing designed primers/probe sets and 25/50 ng input of bisulfite-converted DNA in a PCR Master Mix, and under the following conditions: 95.0°C for 5 minutes followed by 45 cycles of 95.0°C for 15 seconds and 63.0°C for 50 seconds. Fluorescence data was collected at the end of each annealing/extension step for determination of the Cq values. The multiplex was performed on the CFX96 7 Real-Time PCR System using in vitro enzymatically methylated human genomic DNA as calibrator. Cq values were measured at fixed thresholds for fluorescence. For each sample, the Cq values of the marker reactions were normalized for DNA concentration using the ACTB Cq value and the ΔΔCq ratio (2^-ΔΔCq^ x 100) was calculated by comparing the marker Cq values to the Cq values of ACTB and calibrator.

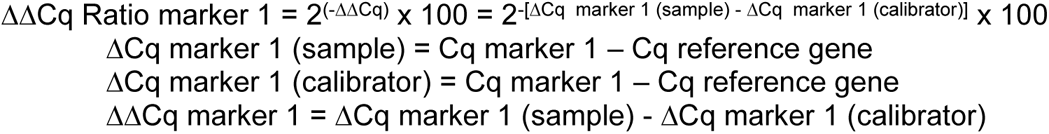

### qMSP data analysis

In the EVAH study there are very few cancer samples as with many other similar studies, this limits the comparison of CIN lesions with cancers. We added our biopsy cancer sample qMSP values to the EVAH study for a better comparison. To compare the similarity of CIN lesions to either healthy control or cancer swabs and cancer biopsy we calculated two different z-scores per biomarker. First, we compared CIN lesions and cancer samples with control samples by taking the mean and standard deviation from the control distribution and using these to calculate a z-score relative to the control samples. Second, we calculated in the same manner a z-score relative to the cancer swab and biopsy samples, using the mean and standard deviation of the cancer distribution. Two thresholds were determined per biomarker cancer threshold; between negative and CIN lesions versus cancer swab and biopsy and CIN3+ threshold; negative, CIN1 and CIN2 versus CIN3, cancer swab and biopsy samples. All values were converted to a z-score for visualization. To summarize four biomarkers results in one scatter plot the average z-score for cancer (y-axis) and control distribution was taken (x-axis). A sample was categorized in low risk if it was not above any threshold, medium risk if above any CIN3+ threshold and below all cancer thresholds and high risk is above any cancer threshold.

---

**Supplemental Figure 1.**
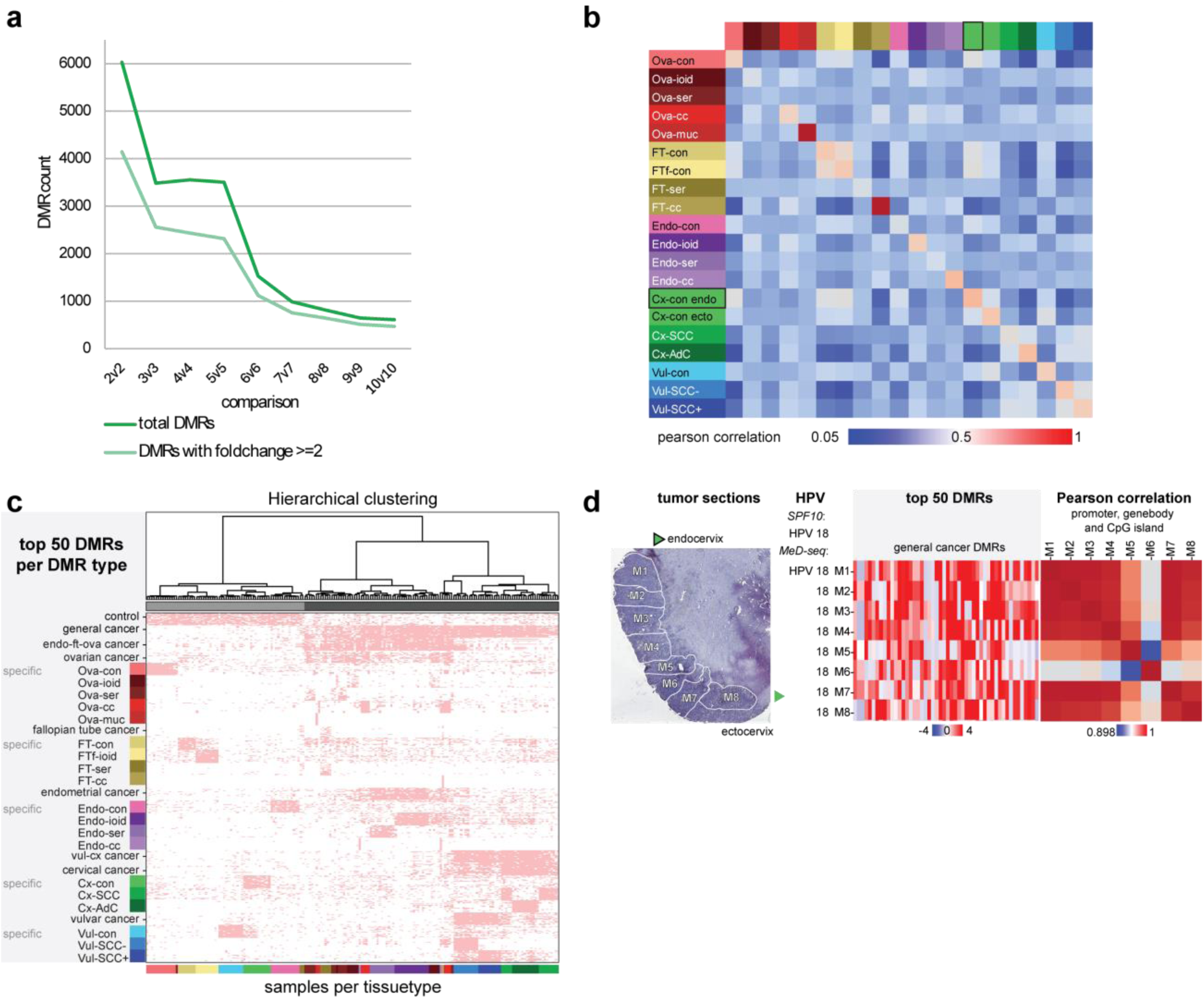
MeD-seq sample size and DMR characteristics. (a) DMR and DMR foldchange>=2 count after DMR analysis using LCM cervix-SCC samples vs cervix-control samples. (b) Average of Pearson correlation between individual samples. (same regions as in figure 2b/2d). (c) Hierarchical clustered heatmap of the top 50 DMRs per DMR type. Sample and DMR type colors are identical to figure 2a. (d) Eight LCM tumor sections with anatomical relation to endo- and ectocervix (left panel). MeD-seq HPV genotype detection per tumor section (middle left panel). Z-score heatmap of the top 50 general cancer markers in different sections (middle right panel), compared to cervix control samples. Pearson correlation of MeD-seq results (right panel) obtained with different tumor sections within the tumor, using promoter, gene body and CpG-island DNA methylation.

**Supplemental Figure 2.**
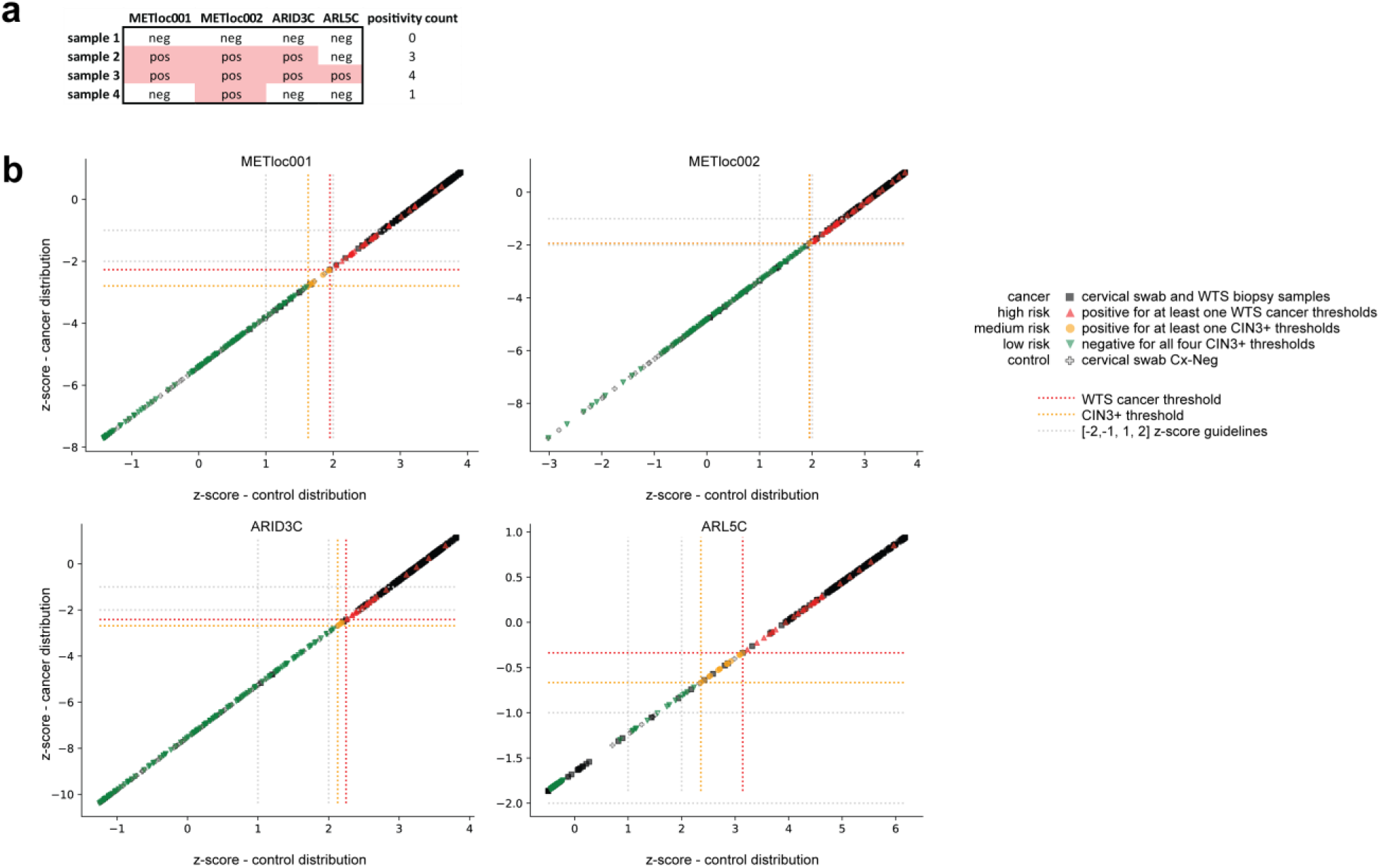
Threshold overview based on control and cancer distribution. (a) Explanation of positivity count per sample based on four biomarkers. (b) Scatter plot per biomarker, z-score based on μ and σ of swab and biopsy control (x-axis) or cancer (y-axis) samples. CIN3+(orange) and cancer (red) threshold show as dotted lines, gray dotted guidelines for z-score of [-2,-1, 1, 2]. CIN lesions subdivided into low, medium and high risk.

